# Diversity and dynamics of fungi during spontaneous fermentations and association with unique aroma profiles in wine

**DOI:** 10.1101/2020.09.10.290536

**Authors:** Di Liu, Jean-Luc Legras, Pangzhen Zhang, Deli Chen, Kate Howell

## Abstract

Microbial activity is an integral part of an agricultural ecosystem and influences the quality of agricultural commodities. Microbial ecology influences grapevine health and crop production, conversion of sugar to ethanol during fermentation, thus wine aroma and flavour. There are regionally differentiated microbial patterns in grapevines and must but how microbial patterns contribute to wine regional distinctiveness (*terroir*) at small scale (<100 km) is not well defined. Here we characterise fungal communities, yeast populations, and *Saccharomyces cerevisiae* populations during spontaneous fermentation using metagenomics and population genetics to investigate microbial distribution and fungal contributions to the resultant wine. We found differentiation of fungi, yeasts, and *S. cerevisiae* between geographic origins (estate/vineyard), with influences from the grape variety. Growth and dominance of *S. cerevisiae* during fermentation reshaped the fungal community and showed geographic structure at the strain level. Associations between fungal microbiota diversity and wine chemicals suggest that *S. cerevisiae* plays a primary role in determining wine aroma profiles at a sub-regional scale. The geographic distribution at scales of less than 12 km supports that differential microbial communities, including the dominant fermentative yeast *S. cerevisiae* can be distinct in a local setting. These findings provide further evidence for microbial contributions to wine *terroir*, and perspectives for sustainable agricultural practices to maintain microbial diversity and optimise fermentation function to craft beverage quality.

## 1. Introduction

Wine grapes (*Vitis vinifera*) are an economically and culturally important agricultural commodity for which microbial activity plays key roles in grape and wine production and quality (Barata et al., 2012; Swiegers et al., 2005). The grapevine harbours complex and diverse microbiota, such as bacteria, filamentous fungi, and yeasts (Barata et al., 2012; Liu and Howell, 2020; Stefanini and Cavalieri, 2018), which substantially modulate vine health, growth, and crop productivity (Berg et al., 2014; Gilbert et al., 2014; Müller et al., 2016). Grapevine-associated microbiota can be transferred to the grape must/juice and have an influence on wine composition, aroma, flavour, and quality (Barata et al., 2012; Ciani et al., 2010; Morrison-Whittle and Goddard, 2018). Wine fermentation is a complex and multispecies process, involving numerous transformations by fungi and bacteria to sculpt chemical and sensory properties of the resulting wines (Swiegers et al., 2005; Verginer et al., 2010). While these consortia all contribute to wine flavour formation, the fermentation process is principally driven by diverse populations of *Saccharomyces cerevisiae* (Fleet, 2003; Goddard, 2008; Howell et al., 2006).

Microbial biogeography contributes to regional distinctiveness of agricultural products, known as “*terroir*” in viticulture [reviewed by Liu et al. (2019)]. Biogeographical patterns in the microbiota associated with the grape and must have been demonstrated for both fungi and bacteria at a regional scale (Bokulich et al., 2014; Mezzasalma et al., 2018; Pinto et al., 2015; Taylor et al., 2014), which are conditioned by multiple factors, such as cultivar, climate and vintage, topography, and soil properties (Bokulich et al., 2014; Liu et al., 2019; Miura et al., 2017; Portillo et al., 2016; Zarraonaindia et al., 2015). Bokulich et al. (2016) showed that the bacterial and fungal consortia correlated with metabolites in finished wines, highlighting the importance of fermentative yeasts (for example, *S. cerevisiae, Hanseniaspora uvarum, Pichia guilliermondii*) and lactic acid bacteria (*Leuconostocaceae*) on the abundance of regional aromatic signatures. Our previous research revealed that wine-related fungal communities structured and distinguished vineyard ecosystems by impacting the flavour and quality of wine, and weather with a contribution by soil properties affected soil and must fungal communities and thus the composition of wines across six winegrowing regions in southern Australia (Liu et al., 2020). However, whether grape-associated microbiota exhibit distinct patterns of distribution at smaller geographic scales (for example individual vineyards) and their associations with wine aroma profiles are not well understood.

Geographic differentiation of *S. cerevisiae* populations is evident at global (Legras et al., 2007; Liti et al., 2009) and regional scales (Gayevskiy and Goddard, 2012; Knight and Goddard, 2015), revealing a picture of distinctive populations at large scales more than ∼ 100 km. Drumonde-Neves et al. (2018) showed higher genetic divergence among *S. cerevisiae* populations between rather than within islands/regions (∼ 1.5 – 260 km scale) and suggested a prevailing role of geography over ecology (grape varieties and agricultural cultivation) in shaping diversification, as previously reported (Goddard et al., 2010). At small scales, several studies characterised significant genetic differences between *S. cerevisiae* populations residing in different vineyards within the same region [Börlin et al. (2016); <10 km] and different sites within a vineyard [Schuller and Casal (2007); 10 – 400 m]. Knight et al. (2015) experimentally demonstrated that regional strains of *S. cerevisiae* produce distinct wine chemical compositions, suggesting a prominent route by which regional *S. cerevisiae* shape wine *terroir*. While few studies have investigated how *S. cerevisiae* differentiation can affect wine aroma, flavour, and characteristics, and none have considered *S. cerevisiae*, fermentative yeasts, and the global fungal communities simultaneously to quantify their contributions to the resultant wine.

To investigate these questions, we sampled microbial communities associated with Pinot Noir and Chardonnay grape must and juice from three wine estates with 8 - 12 km pairwise distances to include grapes from 11 vineyards in the Mornington Peninsula wine region of Victoria, Australia. Using culture-independent sequencing to characterise fungal communities, we disentangled the influences of geographic origin (estate/vineyard), grape variety, and (spontaneous) fermentation stage on the diversity, structure, and composition of the fungal communities. Yeast populations were isolated during spontaneous wine fermentation and taxonomically identified, and the *S. cerevisiae* populations differentiated using microsatellite analysis. To identify the volatiles that differentiated the wine estates we used headspace solid-phase microextraction gas-chromatographic mass-spectrometric (HS-SPME– GC-MS) for metabolite profiling of the resultant wines. Associations between fungal communities and wine metabolites were elucidated with partial least squares regression (PLSR) and structural equation model (SEM). We demonstrate that the grape/wine microbiota and metabolites are geographically distinct, identify multiple layers of fungal microbiota that correlate with wine aroma profiles, and demonstrate that distinctive *S. cerevisiae* exert the most powerful influences on wine quality and style at small geographic scales.

## 2. Materials and methods

### 2.1 Sampling

Five *Vitis vinifera* cv. Pinot Noir and six *Vitis vinifera* cv. Chardonnay vineyards from three wine estates (designated A, B, and C) in the Mornington Peninsula region were selected to conduct this study in 2019 (Supplementary Fig. S1). The distance between wine estates A and B, A and C, B and C is 8 km, 12 km, and 10 km, respectively. Within these estates, vineyards are within a 5 km radius of one another. All vineyards were commercially managed using similar viticultural practices, for example, grapevines were under vertical shoot positioning trellising systems and were applied with the same sprays. Chemical constituents of harvested grapes (°Brix, pH, total acidity) were similar and listed in Supplementary Table S1. Fermentations sampled in this study were conducted without addition of commercial yeasts following similar fermentation protocols. Tanks were cleaned and decontaminated before filling grapes. For Pinot Noir, crushed grapes were held for three days at a cool temperature (known as cold-soaking) and followed by warming so fermentation could commence. Fermentation samples were collected at three time points in duplicate: before fermentation (BF; destemmed and crushed grape musts of Pinot Noir, Chardonnay juice following clarification), at the middle of fermentation (MF, around 50% of sugar fermented), and at the end of fermentation (EF, before pressing, 6-7 °Brix) (Supplementary Table S1). Samples (n = 66) were shipped on ice to the laboratory. Each sample was divided into two subsamples, one was used immediately to isolate yeasts and the other was stored at -20°C for DNA extraction, next -generation sequencing and wine volatile analysis.

### 2.2 Wine volatile analysis

Volatile compounds of EF samples were determined using headspace solid-phase microextraction gas-chromatographic mass-spectrometric (HS-SPME–GC-MS) method (Liu et al., 2016; Zhang et al., 2015) with some modifications. Analyses were conducted with Agilent 6850 GC system and a 5973 mass detector (Agilent Technologies, Santa Clara, CA, USA), equipped with a PAL RSI 120 autosampler (CTC Analytics AG, Switzerland). In brief, 10 mL wine sample was added to a 20 ml glass vial containing 2 g sodium chloride and 20 μL internal standard (4-Octanol, 100 mg/L), and then equilibrated at 35 °C for 15 min. A polydimethylsiloxane/divinylbenzene (PDMS/DVB, Supelco) 65 μm SPME fibre was immersed in the headspace for 10 min at 35°C with agitation, and followed by desorbing in the GC injector for 4 min at 220 °C. Volatile compounds were separated on an Agilent J&W DB-Wax Ultra Inert capillary GC column (30 m × 0.25 mm × 0.25 μm), with helium carrier gas at a flow rate of 0.7 mL/min. The column temperature program was as follows: holding 40 °C for 10 min, increasing at 3.0 °C/min to 220 °C and holding at this temperature for 10 min. The temperature of the transfer line of GC and MS was set at 240 °C. The ion source temperature was 230 °C. The MS was operated in positive electron ionization (EI) mode with scanning over a mass acquisition range of 35 to 350 m/z. Raw data were processed with Agilent ChemStation Software for qualification and quantification. Volatile compounds (n = 79) were identified in wine samples according to retention indices referencing standards and mass spectra matching with NIST11 library. 13 successive levels of standards in model wine solutions (12% v/v ethanol saturated with potassium hydrogen tartrate and adjusted to pH 3.5 using 40% w/v tartaric acid) were analysed by the same protocol as wine samples to establish the calibration curves for quantification. Peak areas of compounds were integrated via target ions model. The concentrations of compounds were calculated with the calibration curves and used for downstream data analysis.

### 2.3 DNA extraction and sequencing

Samples were thawed, and biomass was recovered by centrifugation at 4,000 × g for 15 min, washed three times in ice-cold phosphate buffered saline (PBS) with 1% polyvinylpolypyrrolidone (PVPP) and centrifuged at 10,000 × g for 10 min (Bokulich et al., 2014). The obtained pellets were used for DNA extraction using PowerSoil™DNA Isolation kits (QIAgen, CA, USA). DNA extracts were stored at -20 °C until further analysis.

Genomic DNA was submitted to Australian Genome Research Facility (AGRF) for amplicon sequencing. To analyse the fungal communities, partial fungal internal transcribed spacer (ITS) region was amplified using the universal primer pairs ITS1F/2 (Gardes and Bruns, 1993). The primary PCR reactions contained 10 ng DNA template, 2× AmpliTaq Gold® 360 Master Mix (Life Technologies, Australia), 5 pmol of each primer. A secondary PCR to index the amplicons was performed with TaKaRa Taq DNA Polymerase (Clontech). Amplification was conducted as follows: 95 °C for 7 min, followed by 35 cycles of 94 °C for 30 s, 55 °C for 45 s, 72 °C for 60 s, and a final extension at 72 °C for 7 min. The resulting amplicons were cleaned again using magnetic beads, quantified by fluorometry (Promega Quantifluor), and normalised. The equimolar pool was cleaned a final time using magnetic beads to concentrate the pool and measured using a High-Sensitivity D1000 Tape on an Agilent 2200 TapeStation. The pool was diluted to 5nM and molarity was confirmed again with a High-Sensitivity D1000 Tape. This was followed by 300 bp paired-end sequencing on an Illumina MiSeq (San Diego, CA, USA).

Raw sequences were processed using QIIME v1.9.2 (Caporaso et al., 2010). Low quality regions (Q < 20) were trimmed from the 5′ end of the sequences, and the paired ends were joined using FLASH (Magoč and Salzberg, 2011). Primers were trimmed and a further round of quality control was conducted to discard full length duplicate sequences, short sequences (< 100 nt), and sequences with ambiguous bases. Sequences were clustered followed by chimera checking using UCHIME algorithm from USEARCH v7.1.1090 (Edgar et al., 2011). Operational taxonomic units (OTUs) were assigned using UCLUST open-reference OTU-picking workflow with a threshold of 97% pairwise identity (Edgar, 2010). Singletons or unique reads in the resultant data set were discarded. Taxonomy was assigned to OTUs in QIIME using the UNITE fungal ITS database (v7.2) (Kõljalg et al., 2005). To avoid/reduce biases generated by varying sequencing depth, sequences were rarefied to 10,000 per sample prior to downstream analysis. Raw sequencing reads are publicly available in the National Centre for Biotechnology Information (NCBI) Sequence Read Archive under the bioproject PRJNA646865.

### 2.4 Yeast isolation and identification

Yeast populations were evaluated during spontaneous fermentations (BF, MF, EF). Aliquots of 0.1 ml from serially diluted samples (from 10^−2^ to 10^−5^) were spread on Wallerstein Laboratory Nutrient (WLN) agar plates (Oxoid, Australia) that were supplemented with 34 mg/mL chloramphenicol and 25 mg/mL ampicillin to inhibit bacterial growth. After 5 days of incubation at 28°C, colonies were counted (30-300 colonies) and colony morphology was recorded to differentiate yeast species according to Pallmann et al. (2001) and Romancino et al. (2008). Characteristic yeast colonies were isolated by subculturing on fresh WLN plates. DNA was extracted from pure colonies using the MasterPure™Yeast DNA Purification Kit (Epicentre, Madison, WI) following the manufacturer’s instructions. The 26S rRNA D1/D2 domain was amplified using primers NL1/4 (Kurtzman and Robnett, 1998) for sequencing by AGRF. Species identity was determined using BLAST hosted by the NCBI (http://blast.ncbi.nlm.nih.gov/Blast.cgi), considering an identity threshold of at least 98%. The colonies identified as *S. cerevisiae* were analysed by microsatellites as described in the following paper. Sequence data was uploaded to Genbank with accession numbers MT821080-MT821098.

### 2.5 Microsatellite characterisation

A set of 94 *S. cerevisiae* strains, listed in Supplementary Table S1, was characterised for allelic variation at 12 microsatellites as previously described (Legras et al., 2007). Briefly, two multiplex assays of primers corresponding to loci C5, C3, C8, C11, SCYOR267c and C9, YKL172w, ScAAT1, C4, SCAAT5, C6, YPL009c, were amplified using the QIAGEN Multiplex PCR Kit according to the manufacturer’s instructions. PCRs were run in a final volume of 12.5 μL containing 10 – 250 ng yeast DNA, and used the following program: initial denaturation at 95°C for 15 min, followed by 34 cycles of 94°C for 30 s, 57°C for 2 min, 72°C for 1 min, and a final extension at 60°C for 30 min. PCR products were sized for 12 microsatellite loci on an ABI 310 DNA sequencer (Applied Biosystems) using the size standards HD400ROX. Allele distribution into classes was carried out using Genious software v9.1.6 (Biomatters) and the corresponding alleles classes were described in Legras et al. 2007.

### 2.6 Data analysis

Alpha diversities of fungal communities, yeast populations, and wine volatile compounds were calculated using the Shannon index with the “vegan” package (Oksanen et al., 2007). One-way analysis of variance (ANOVA) was used to determine whether sample classifications (e.g., fermentation stage, wine estate) contained statistically significant differences in the alpha-diversity. Principal coordinate analysis (PCoA) was performed to evaluate the distribution patterns of fungal communities, yeast populations, and wine aroma based on beta-diversity calculated by the Bray– Curtis distance with the “labdsv” package (Roberts, 2007). Permutational multivariate analysis of variance (PERMANOVA) using distance matrices with 999 permutations was conducted within each sample class to determine the statistically significant differences with “adonis” function in “vegan” (Oksanen et al., 2007). Significant taxonomic differences of fungi in the BF must between sample categories (wine estate, grape variety) were tested using linear discriminant analysis (LDA) effect size (LEfSe) analysis (Segata et al., 2011) (https://huttenhower.sph.harvard.edu/galaxy/). The OTU table was filtered to include only OTUs > 0.01% relative abundance to reduce LEfSe complexity. The factorial Kruskal–Wallis sum-rank test (*α* = 0.05) was applied to identify taxa with significant differential abundances between groups (all-against-all comparisons), followed by the logarithmic LDA score (threshold = 2.0) to estimate the effect size of each discriminative feature. Significant taxa were used to generate taxonomic cladograms illustrating differences between sample categories. Dynamics and successions of the dominant species (relative abundance > 1.00%) demonstrating significant differences among stages (ANOVA; FDR-corrected *p* value < 0.05) were illustrated by alluvial diagrams using the “ggalluvial” package (Brunson, 2018) in “ggplot2”.

Multilocus genotypes were calculated with the “poppr” package (Kamvar et al., 2014). The Bruvo’s distance (Bruvo et al., 2004) was calculated between each strain by the “ape” (Paradis et al., 2004) and “poppr” packages. The tree was obtained from the distance matrices with “poppr”, and drawn using MEGA X (Kumar et al., 2018). The tree was rooted by the midpoint method. An alternative method of genetic clustering-discriminant analysis of principal components (DAPC) was also applied to infer the population structure with “adegenet” package (Jombart et al., 2010). Analysis of molecular variance (AMOVA) was conducted to determine the degree of differentiation of variations between designated partitions (estates, vineyards, varieties) with “poppr” (Excoffier et al., 1992). The genetic distance *F*_*ST*_(Reynolds et al., 1983) between estates were calculated by the “hierfstat” package (Goudet, 2005). PCoA based on the Bruvo’s distance was conducted to evaluate the geographic distribution pattern of *S. cerevisiae* populations with “vegan” (Oksanen et al., 2007).

Partial least squares regression (PLSR) analysis with cross-validation was used to model associations between normalised mean values for fungal taxa (species with relative abundance > 0.10%) and wine volatile compounds using the “pls” package (Wehrens and Mevik, 2007). The structural equation model (SEM) (Grace, 2006) was used to evaluate the direct and indirect relationships between must fungal communities and yeast populations, *S. cerevisiae* populations, and resulting wine aroma (the first axis values of PCoA analysis). SEM is an a priori approach partitioning the influence of multiple drivers in a system to help characterise and comprehend complex networks of ecological interactions (Eisenhauer et al., 2015). An *a priori* model was established based on the known effects and relationships among drivers of distribution patterns of wine aroma to manipulate the data before modelling (including must aroma, °Brix, pH; data not shown). A path coefficient described the strength and sign of the relationship between two variables (Grace, 2006). The good fit of the model was validated by the χ2-test (*p* > 0.05), using the goodness of fit index (GFI > 0.90) and the root MSE of approximation (RMSEA < 0.05) (Schermelleh-Engel et al., 2003). Standardised total effects of each factor on the wine aroma distribution pattern were calculated by summing all direct and indirect pathways between two variables (Grace, 2006). All these analyses were conducted using AMOS v25.0 (AMOS IBM, NY, USA).

## 3. Results

### 3.1 Fungal microbiota vary by geographical origin and grape variety

To elucidate the influences of geographic locations, grape variety, and fermentation process on the wine microbiota, 66 duplicate samples covering three wine estates, Pinot noir and Chardonnay, from the beginning, middle and end of fermentation were collected to analyse fungal communities. A total of 1,566,576 ITS high-quality sequences were generated from all samples, which were clustered into 277 fungal OTUs with a threshold of 97% pairwise identity. *Ascomycota* was the most abundant phylum in the grape must/juice before fermentation (BF) comprising 92.00% of all sequences, followed by *Basidiomycota* (7.98%) and *Mortierellomycota* (< 0.01%) (Data not shown). Fungal profiles were dominated by filamentous fungi, mostly of the genera *Cladosporium, Aureobasidium*, and *Epicoccum*, with notable populations of fermentative yeasts including *Saccharomyces* and *Hanseniaspora*, as well as the basidiomycetous yeasts *Vishniacozyma, Rhodotorula*, and *Sporobolomyces* (Fig. 1A).

**Fig. 1.**
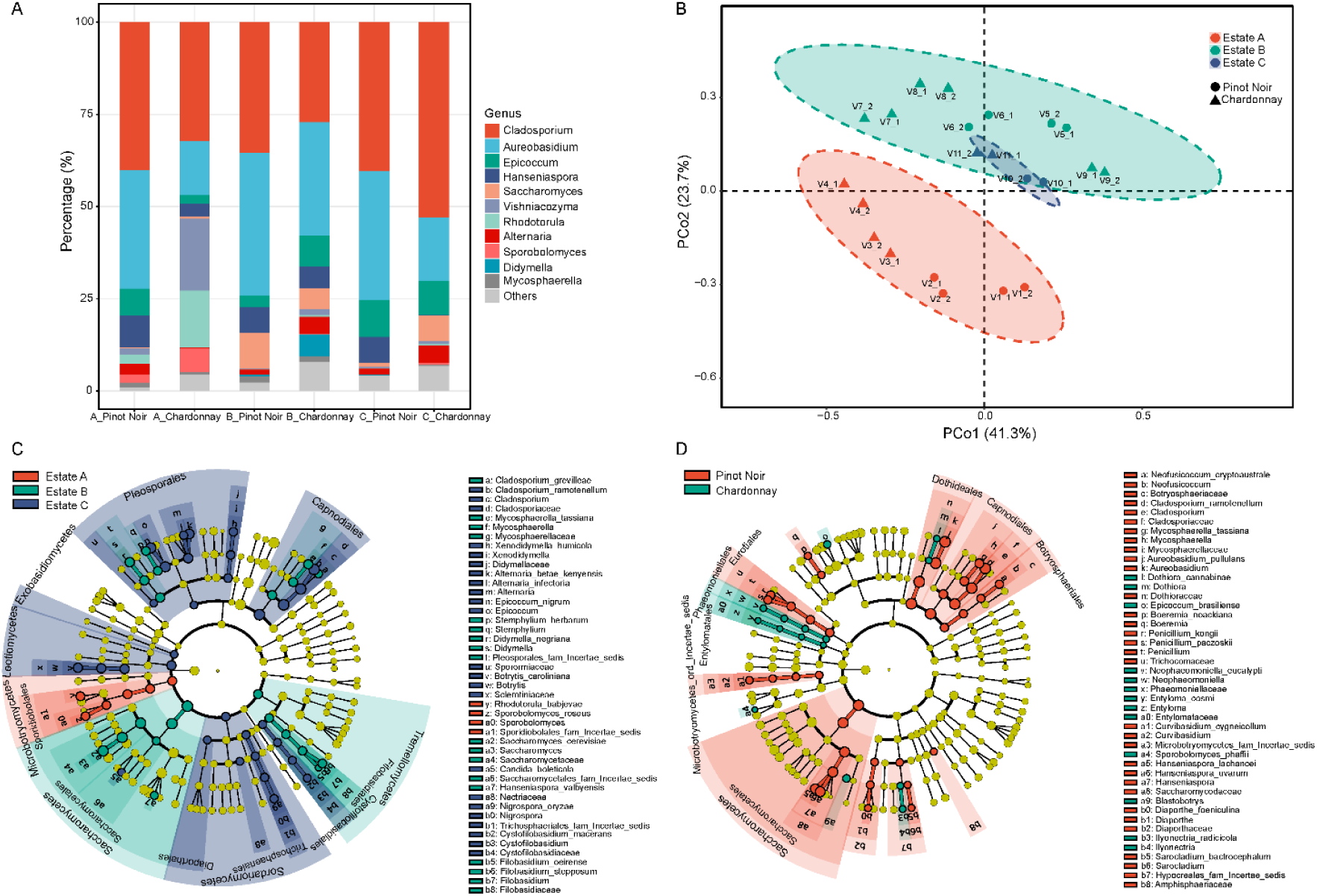
Musts before fermentation (BF) have differential fungal communities depending geographical origins and grape varieties. (A) Microbial community composition characterised to the genus level (11 genera, relative abundance > 1.00% shown); (B) Principal coordinate analysis (PCoA) based on Bray-Curtis distances among wine estates across both varieties; (C) Linear discriminant analysis (LDA) effect size (LEfSe) taxonomic cladogram identifying significantly discriminant (Kruskal–Wallis sum-rank test *α* < 0.05; LDA score > 2.00) taxa associated with wine estates; (D) LEfSe taxonomic cladogram identifying significantly discriminant taxa associated with grape varieties.

Individual vineyards and wine estates were distinguished based on the microbial community present in the grape must/juice before fermentation (Fig. 1B; Supplementary Table S2). Permutational multivariate analysis of variance (PERMANOVA) based on the Bray–Curtis distance confirmed that fungal composition was significantly different between estates (R^2^ = 0.310, *p* < 0.001) and at least two vineyards (R^2^ = 0.573, *p* < 0.001) regardless of grape variety. Pinot Noir demonstrated stronger geographical differentiation (PERMANOVA, R^2^_Estate_ = 0.683, *p* < 0.001; R^2^_Vineyard_= 0.779, *p* < 0.001) than Chardonnay (PERMANOVA, R^2^_Estate_ = 0.470, *p* < 0.001; R^2^_Vineyard_= 0.566, *p* < 0.001). Grape variety weakly but significantly impacted fungal composition (PERMANOVA, R^2^ = 0.103, *p* = 0.021). This impact was more distinct with an improved coefficient of determination (R^2^) within a certain wine estate (Fig. 1B; Supplementary Table S2). Principal coordinate analysis (PCoA) showed the geographic patterns (95% confidence interval based on estates) of fungal communities across both varieties, with 65.0% of total variance explained by the first two principal coordinate (PC) axes (Fig. 1B). Linear discriminant analysis (LDA) effect size (LEfSe) analysis further confirmed that these patterns related to significant associations (Kruskal–Wallis sum-rank test, *α* < 0.05) between fungal taxa and wine estates (Fig. 1C), and grape varieties (Fig. 1D), respectively. *Microbotryomycetes*, notably *Rhodotorula babjevae* and *Sporobolomyces roseus*, were observed with higher abundances in the grape must/juice from wine estate A, with *Saccharomycetaceae* (notably fermentative yeast *S. cerevisiae*), *Tremellomycetes* (including *Filobasidium* spp.), *Cladosporium grevilleae, Mycosphaerellaceae, Stemphylium, Didymella* from wine estate B, and *Capnodiales* (notably *Cladosporium* spp.), *Pleosporales* (including *Didymellaceae, Alternaria, Epicoccum, Sporormiaceae*), *Exobasidiomycetes, Leotiomycetes* (including *Botrytis*), *Sordariomycetes* (including *Cystofilobasidiales, Trichosphaeriales, Diaporthales*) from wine estate C. For varietal influences, *Phaeomoniellales* including *Phaeomoniellaceae* and *Entylomataceae, Dothiora, Epicoccum brasiliense, Sporobolomyces phaffii, Blastobotrys*, and *Ilyonectria* were significantly abundant in Chardonnay juice, while *Botryosphaeriales, Capnodiales* (in particular *Cladosporium ramotenellum*), *Dothideales* (in particular *Aureobasidium pullulans*), *Eurotiales* (including *Penicillium*), *Saccharomycetes* (in particular *Hanseniaspora* spp.), *Microbotryomycetes_ord_Incertae_sedis, Boeremia, Diaporthaceae, Sarocladium*, and *Amphisphaeriaceae* in Pinot Noir musts.

### 3.2 Microbiota dynamics during spontaneous wine fermentation

Significant decreases in the fungal diversity (ANOVA, *p* < 0.001) were recorded between fermentation stages regardless of grape variety (Fig. 2A, Supplementary Fig. S2A). PCoA showed that ferments were grouped according to their fermentative stage, where PC1 explained 66.0% and 57.3% of the total variance for Pinot Noir (Fig. 2B; PERMANOVA, R^2^ = 0.648, *p* < 0.001) and Chardonnay (Supplementary Fig. S2B; PERMANOVA, R^2^ = 0.608, *p* < 0.001), respectively. BF samples were clearly separated from MF and EF, while the latter groups were partially overlapped. Within grape varieties, tracking major species across vineyards and estates (relative abundance > 1.00%) revealed fungal dynamics and succession during fermentation. All these species shown in the alluvial diagrams presented significantly different relative abundances as the fermentation progressed (ANOVA; FDR-corrected *p* value < 0.05) (Fig. 2C, Supplementary Fig. S2C). BF musts/juice displayed diverse and variable collections of fungi, of which *A. pullulans* and *C. ramotenellum* were the most abundant species for Pinot Noir and Chardonnay, respectively. In the beginning of Pinot Noir fermentation, filamentous fungi and non-*Saccharomyces* yeasts dominated 89.9% of the community, with 5.45% relative abundance for *S. cerevisiae*. After fermentation started, *S. cerevisiae* grew and gradually dominated the community, occupying 66.5% of the MF community and 80.9% of the EF community, while other species underwent drastic decreases in relative abundances during the fermentation (Fig. 2C). Likewise, this succession pattern was also observed in Chardonnay ferments, except that few taxa (*A. pullulans, R. babjevae, Hanseniaspora lachancei*) showed slight recoveries in relative abundances between MF and EF (Supplementary Fig. S2C). Correspondingly, geographical differences in fungal communities were not significant based on estates in both MF (PERMANOVA, R^2^ = 0.162, *p* = 0.104) and EF (PERMANOVA, R^2^ = 0.145, *p* = 0.162) ferments, although fungi differentiated vineyard origin of some vineyards in both stages (*p* < 0.001) (Supplementary Table S2).

**Fig. 2.**
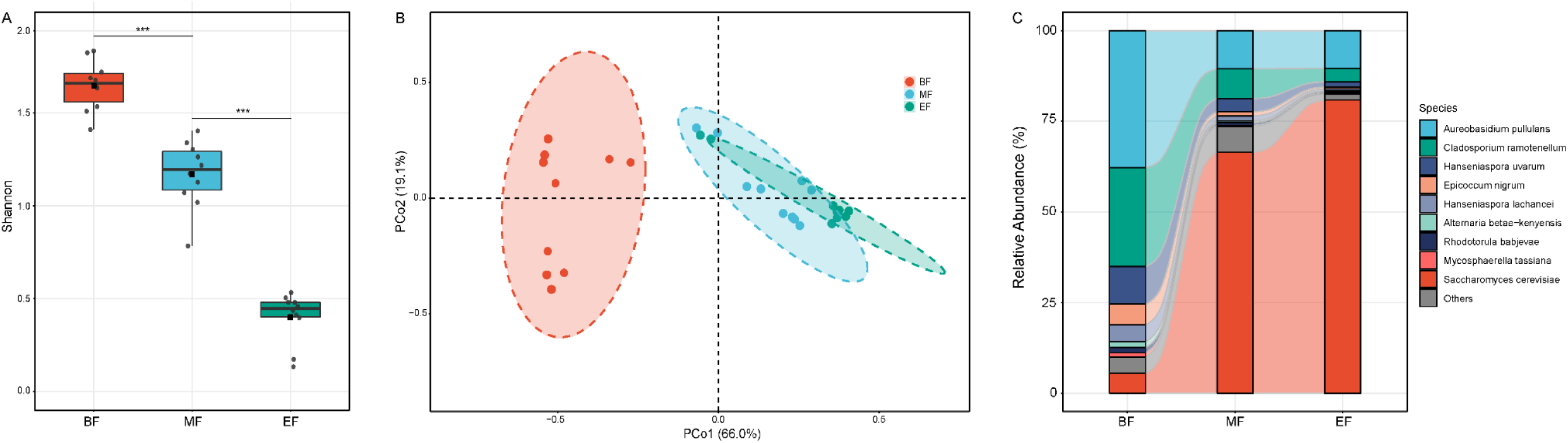
Stage of fermentation influences microbial diversity and composition of Pinot Noir. (A) *α*-diversity (Shannon index) significantly decreases (*p* < 0.001***) during the fermentation; (B) Bray-Curtis distance PCoA of fungal communities according to the fermentation stage (BF, before fermentation; MF, at the middle of fermentation; EF, at the end of fermentation); (C) Relative abundance changes of major fungal species (9 species; relative abundance > 1.00%) that significantly differed by fermentation stage (ANOVA; FDR-corrected *p* value < 0.05).

### 3.3 Yeast population dynamics during spontaneous wine fermentation

To estimate yeast population dynamics during spontaneous fermentations, we isolated colonies based on morphology on WLN medium and identified yeasts using the taxonomically distinctive 26S rRNA D1/D2 region. A total of 359 yeast isolates were obtained (Supplementary Table S1), corresponding to 14 species: *Hanseniaspora uvarum, Hanseniaspora opuntiae, Torulaspora delbrueckii, Metschnikowia andauensis, Meyerozyma guilliermondii, Candida africana, Candida intermedia, Candida oleophila, Candida ishiwadae, Rhodotorula mucilaginosa, Pichia membranifaciens, Wickerhamomyces anomalus, Saccharomyces bayanus*, and *S. cerevisiae* (Supplementary Table S3). All of these species had also been identified with ITS amplicon sequencing (give figure/table number). *S. cerevisiae* and the non-*Saccharomyces* species of *H. uvarum, H. opuntiae, T. delbrueckii, M. andauensis* dominated the isolates, whereas other species appeared sporadically. At the beginning of fermentation, the grape must/juice harboured high species diversity (Shannon index) of yeasts, with non-*Saccharomyces* yeasts dominating the populations (Supplementary Table S3). The amount and distribution of yeast species differed among vineyards, estates, and varieties (Supplementary Table S3), with significant geographical differences observed in population compositions (PERMANOVA based on Bray–Curtis distance, R^2^_Estate_ = 0.287, *p* = 0.019; R^2^_Vineyard_= 0.426, *p* = 0.002). As fermentation proceeded, the viable population of yeast in the must increased from initial values of 10^4^–10^6^ to 10^7^–10^8^ CFU/mL in the middle fermentation (MF) and declined coinciding with decreased species diversity during fermentation (Supplementary Table S3). Non-*Saccharomyces H. uvarum* and *H. opuntiae* were isolated throughout some fermentations, with *T. delbrueckii* isolated at the beginning and middle fermentation points. In spite of low initial abundance, *S. cerevisiae* dominated the ferments at middle and end fermentation points and occupied 100% of some yeast populations from wine estate B (Supplementary Table S3).

From the isolated yeasts, 94 *S. cerevisiae* strains were further analysed through 12 microsatellite loci, resulting in 80 multilocus genotypes, with 14 strains showing genotypes identical to others in this study. The 12 microsatellite loci recorded from 4 to 23 different alleles per locus, of which C5 and SCAAT1 displayed the highest diversity with 24 and 21 alleles, respectively. Two strains were shared between vineyards and varieties, while other identical strains were isolated from within a single vineyard and are likely the same strain. The neighbour-joining tree built from the Bruvo’s distance showed clustering linked to the geographica l origin of strains (Fig. 3A). Some branches clustered isolates from one wine estate with very close genetic relationships as group I (A), group II (B), and group III (C). Some strains from estate A and B gathered in clusters, with only few strains from estate C (group IV, V). Group VI was composed of clusters originating from all three estates. The relationships among *S. cerevisiae* strains inferred using DAPC were consistent with the genetic tree (Fig. 3B). The total amount of genetic variation explained by the first 35 principal components was 93.8%, of which 24.12% was conserved by the first two axes. Most populations were clearly separated into estate A, B, and C, with overlaps observed among groups (Fig. 3B). Strains V1.2.1, V9.1.2, and V11.2.4 among three estates were coinciding with group VI in the phylogenetic tree, as well as strains V4.3.4 and V7.2.6 between estate A and B clusters with group IV (Fig.3). AMOVA confirmed significant influences of geographic origins (estate/vineyard, *p* = 0.001) and the grape variety (*p* = 0.023) on the *S. cerevisiae* population structure based on, in particular based on estates (Supplementary Table S4). Within estates, the difference between vineyards or varieties was not significant (Supplementary Table S4). Higher differentiation between was observed between wine estates A and C (12 km; F_*ST*_ *=* 0.143, *p* < 0.001), B and C (10 km; F_*ST*_ *=* 0.133, *p* < 0.001), than between A and B (8 km; F_*ST*_ *=* 0.076, *p* < 0.001) (Supplementary Table S4).

**Fig. 3.**
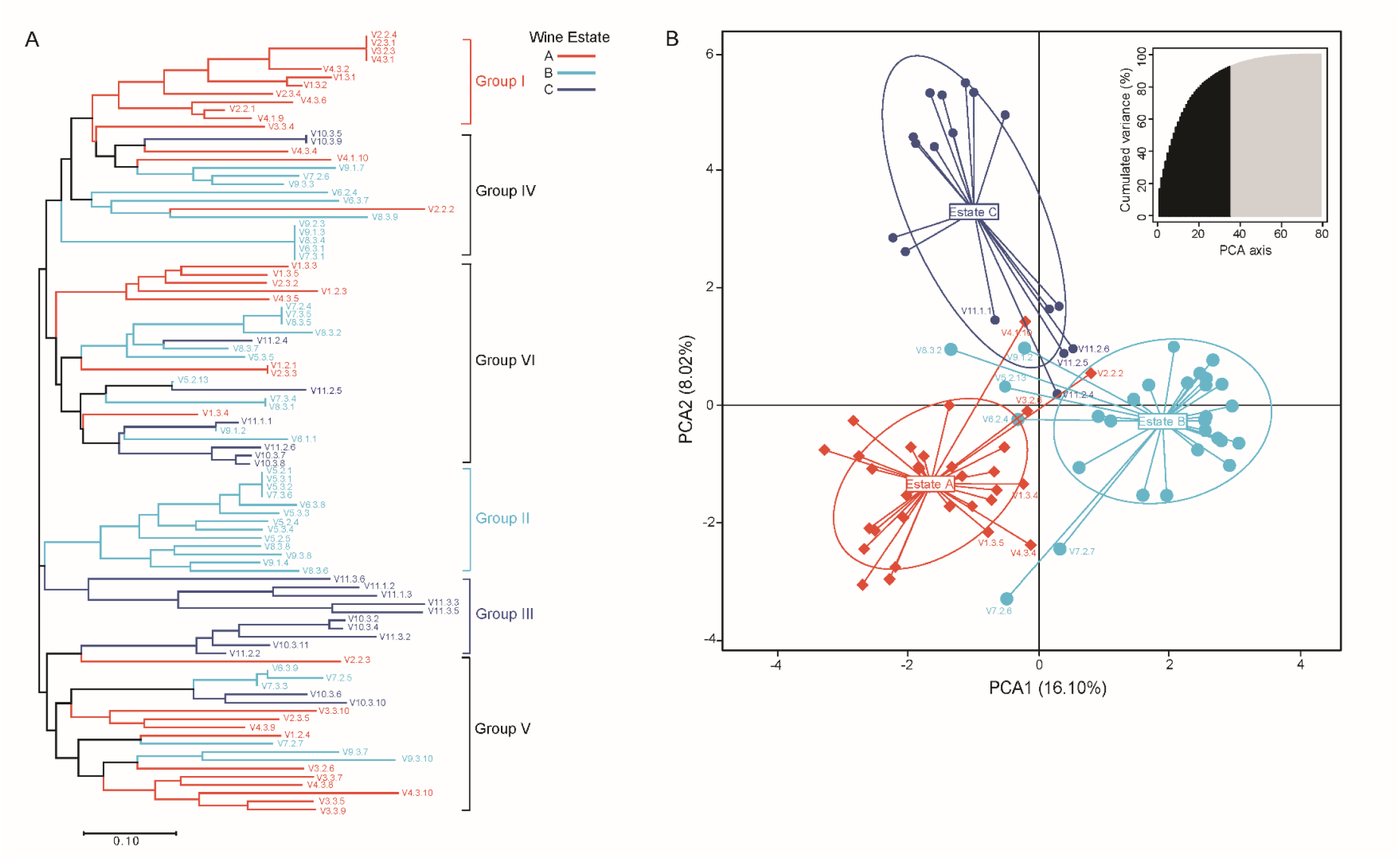
*Saccharomyces cerevisiae* populations show geographic clustering. (A) Neighbour-joining tree showing the clustering of 94 *S. cerevisiae* strains isolated from three wine estates. The tree was constructed from the Bruvo’s distance between strains based on the polymorphism at 12 loci and is rooted according to the midpoint method. Branches are coloured according to the wine estate from which strains have been isolated. (B) Scatterplot from discriminant analysis of principal components (DAPC) of the first two principal components discriminating *S. cerevisiae* populations by estates. Points/diamonds represent individual observations, and lines represent population memberships. Inertia ellipses represent an analog of a 67% confidence interval based on a bivariate normal distribution. Number of principal components (n = 35) at which the maximal reassignment of samples occurred are depicted as black lines the PCA graph on the topright corner, with subsequent components in grey line.

### 3.4 Aroma profiles are distinctive for wines of each geographical origin

Using GC-MS, we analysed the volatile compounds of Pinot Noir and Chardonnay wine samples (at the end of fermentation) in triplicate to represent wine metabolite profiles. In all, 79 volatile compounds were identified in these wines. Pinot Noir wines contained 37 geographically differential compounds based on wine estates, and Chardonnay wines contained 46 (Supplementary Table S5). Within grape varieties, wine complexity (as determined by Shannon index) was not significantly different amongst wine estates (ANOVA; F_Pinot Noir (2, 12)_ = 2.393, *p* = 0.161; F_Chardonnay (2, 15)_ = 2.313, *p* = 0.284). Wine aroma profiles are clearly separated with PCoA based on Bray–Curtis dissimilarity according to estates, where PC1 explained 65.8% and 62.7% of the total variance for Pinot Noir wines (PERMANOVA, R^2^ = 0.722, *p* = 0.011) and Chardonnay wines (PERMANOVA, R^2^ = 0.773, *p* = 0.002), respectively (Fig. 4).

**Fig. 4.**
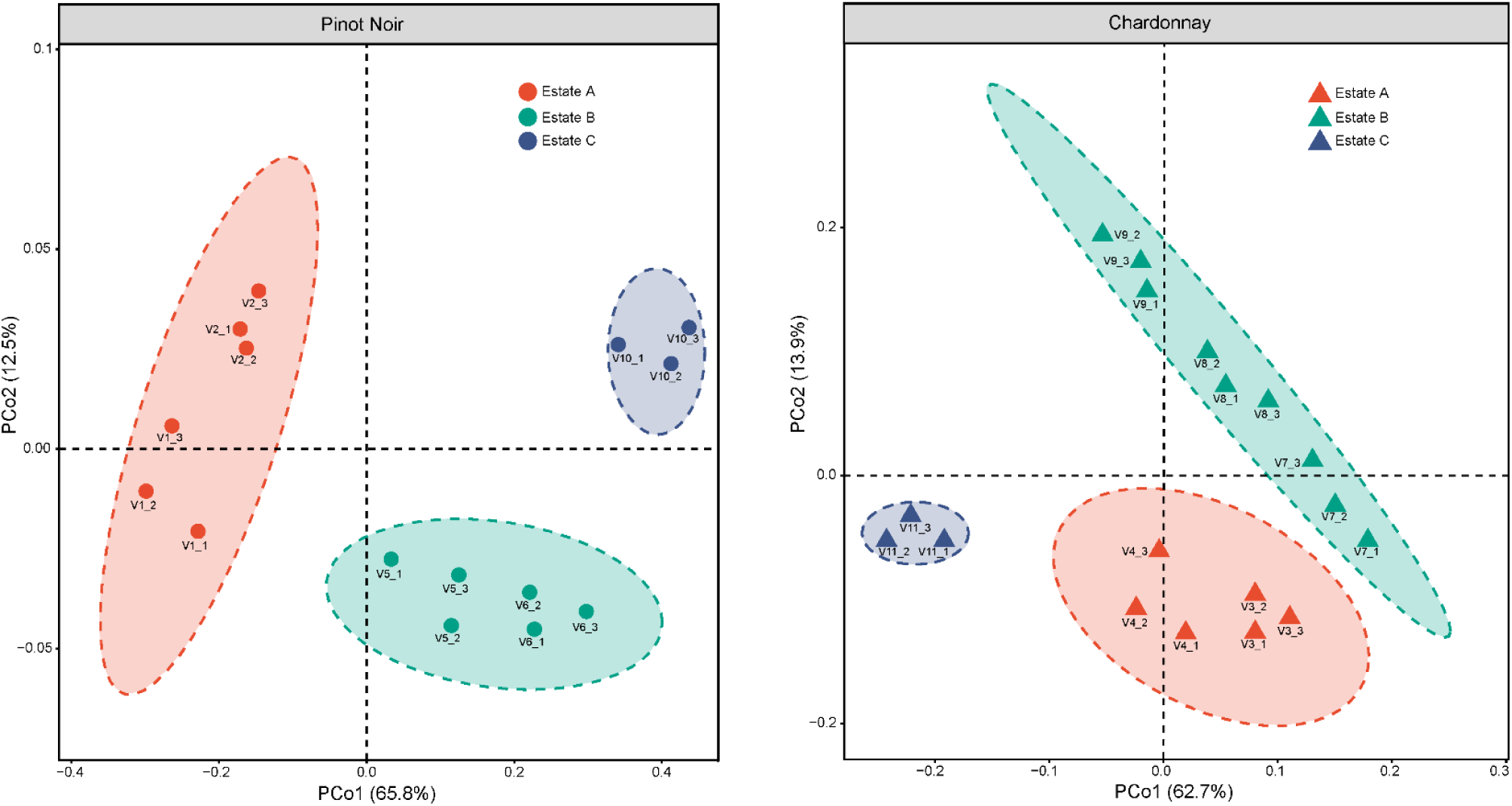
Wine metabolites exhibit geographical variation. PCoA based on Bray-Curtis dissimilarity obtained from comparing volatile profiles of Pinot Noir and Chardonnay wines.

### 3.5 Fungal microbiota correlate to wine aroma profiles

To elucidate the relationship between geographically differential fungal microbiota and wine metabolites, partial least squares regression (PLSR) was used to model covariance between fungal species and volatile compounds. PSLR projections were made with dominant species in the grape must (relative abundance > 0.01% across samples; Pinot Noir, 20 species; Chardonnay, 23 species) and volatile compounds in the resulting wines. Compounds shown in the plots explained > 20% of the variance in the first two components (Fig. 5), of which many were the same compounds identified by ANOVA (Supplementary Table S5, S6). PLSR showed highly covariable relationships between fermentative yeasts and volatile compounds. In Pinot Noir wines, some interesting correlations were observed; for example *S. cerevisiae* and *T. delbrueckii* correlated strongly with several esters (C33, Ethyl octoate; C26, Ethyl lactate; C55, 3-Methylbutyl octanoate; C38, Ethyl 6-heptenoate); *M. guilliermondii* with monoterpenes (C43, linalool; C48, terpinen-4-ol; C68, nerol; C59, *α*-terpineol), *β*-damascenone (C70), nonanals (C41, 2-nonanol; C56, 1-nonanol) and the related ester (C42, ethyl nonanoate); and *H. uvarum* and *H. lachancei* with alcohols (C46, 2,3-butanediol; C63, 1-decanol) and esters (C62, methyl salicylate; C76, ethyl tetradecanoate). In Chardonnay wines, *S. cerevisiae* was associated with some esters (for example, C33; C55; C16, ethyl hexanoate; C61, benzyl acetate; C20, hexyl acetate), *T. delbrueckii* and *M. guilliermondii* with some alcohols (C63; C44, 1-octanol) and esters (C26; C38; C42; C76), and *H. uvarum* and *H. lachancei* with C46, C62, and fatty acids (C45, 2-methyl-propanoic acid; C72, hexanoic acid; C77, octanoic acid).

**Fig. 5.**
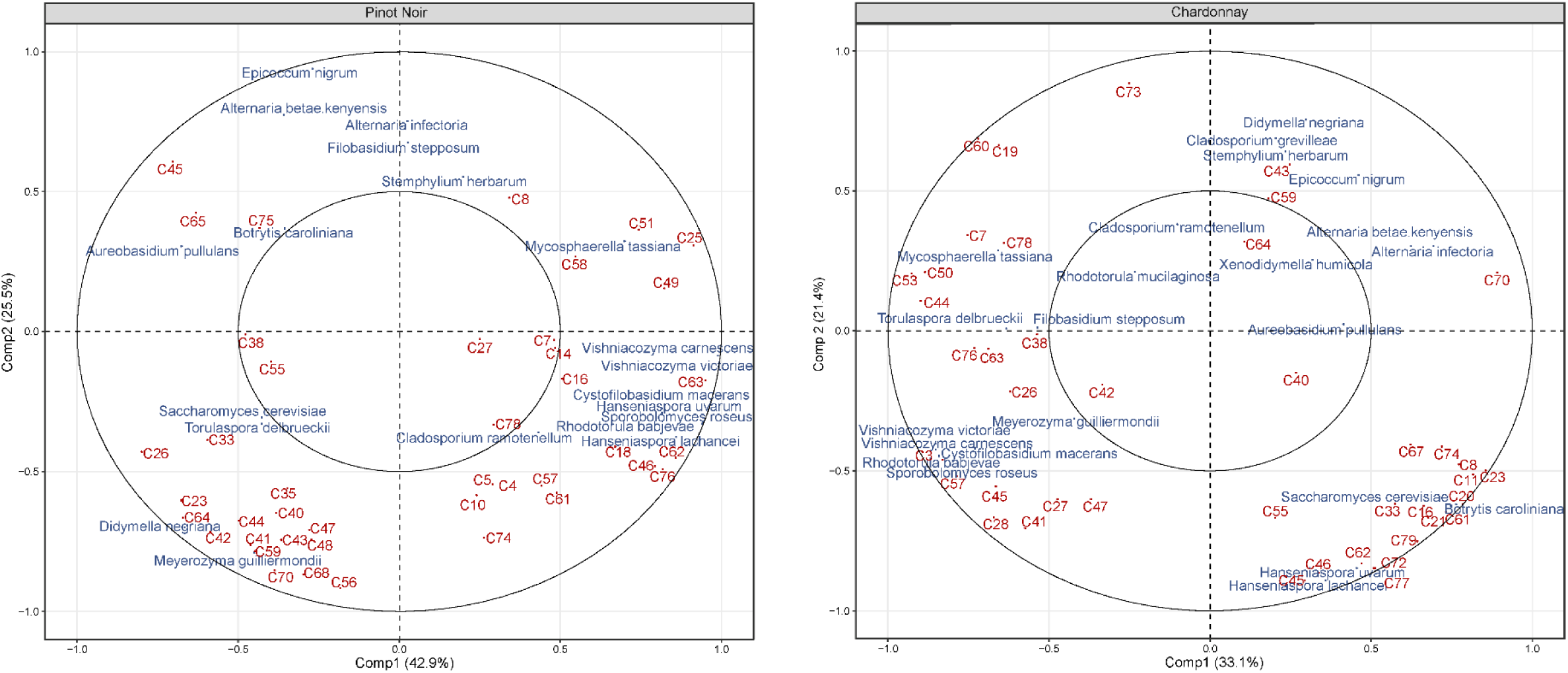
Partial least squares regression (PLSR) demonstrates microbial influence on volatile compounds of Pinot Noir and Chardonnay wines.

To disentangle the role of microbial communities on wine metabolites, we used structural equation modelling (SEM) (Grace, 2006) to testify fungal community compositions (the first axis of PCoA) at multiple levels simultaneously: fungal communities, yeasts (*S. cerevisiae* and non-*Saccharomyces* yeasts), and *S. cerevisiae* populations. The SEM explained 84.9% of the variance found in the geographical pattern of wine aroma (Fig. 6). Fungal communities in the must drove wine aroma profiles directly (path coefficient = 0.286***) and indirectly by effects on yeasts and *S. cerevisiae* populations, in particular strong influences on yeast populations (path coefficient = 0.562). *S. cerevisiae* populations had the highest direct positive effects on the resulting wine aroma characteristics, while yeast populations had the lowest but significant effects (Fig. 6A). Overall, *S. cerevisiae* populations were the most important driver of wine characteristics (Fig. 6B).

**Fig. 6.**
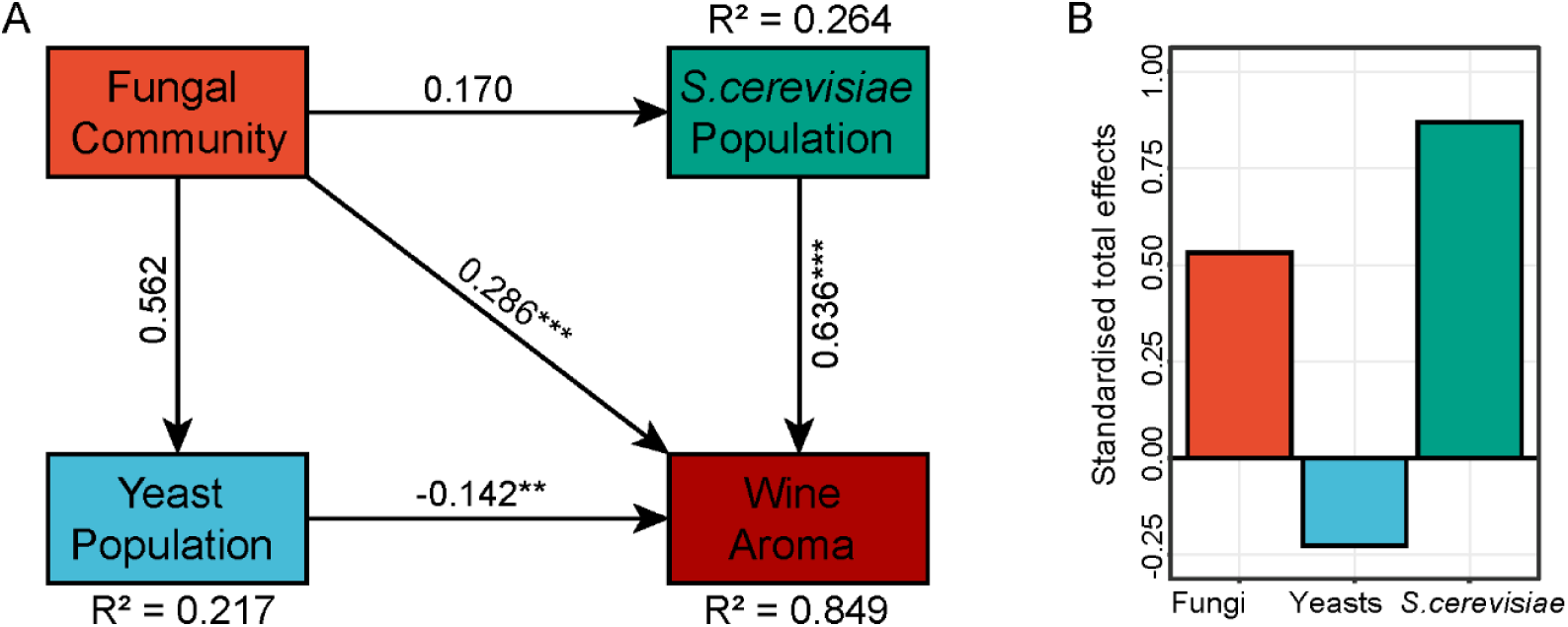
Direct and indirect effects of fungal community composition (the first axis of PCoA) at multiple levels on wine aroma profiles. Structural equation model (SEM) fitted to wine aroma composition (A) and standardised total effects (direct plus indirect effects) derived from the model (B).

## 4. Discussion

There is mounting evidence for geographical differentiation of wine-related microbial communities at regional scales (Bokulich et al., 2014; Gayevskiy and Goddard, 2012; Jara et al., 2016; Pinto et al., 2015; Taylor et al., 2014). Our previous work revealed that different wine-producing regions in southern Australia possess distinct, distinguishable microbial patterns (especially fungal microbiota) at the scale of 400 km, correlated with local weather conditions and soil properties (Liu et al., 2020). In the current study, within a single sub-region spanning 12 km, we demonstrate geographical differentiation of grape must/juice fungal communities, yeast populations, and *S. cerevisiae* populations, with influences from the grape variety.

### 4.1 Fungal ecology at multiple layers during spontaneous wine fermentation

In the freshly crushed grape must/juice, fungal communities were highly diverse and characterised by ubiquitous genera such as *Aureobasidium, Cladosporium, Saccharomyces*, and *Rhodotorula*, deriving from the vineyard ecosystem. Geographical origin had a greater impact on the fungal community (Supplementary Table S2) and yeast populations (results 3.2) than grape variety, and this is in line with other studies in grapevine-associated microbiota (Bokulich et al., 2014; Mezzasalma et al., 2018; Wang et al., 2015). In particular, *S. cerevisiae* yeasts was one of geographical features (Fig. 1C). Given the small scale of the vineyards in this study (< 12 km), macroclimate does not differentiate the sites, and so local conditions appear to modulate communities. Geographical features (for example vineyard orientation), microclimate, and soil properties (for example nutrient availability) could explain some variation among vineyard sites, but these measures were beyond the scope of this study. Within a single estate, grape variety played a significant role in shaping fungal communities (Fig. 1B; Supplementary Table S2), suggesting a genetic component to plant-microbial interactions (Bokulich et al., 2014; Mezzasalma et al., 2018). Cultivar variation in the microhabitat and environmental stress responses may explain how grapevines recruit their associated microbiota, including both the normal microbiota and cultivar-specific susceptibilities to disease pressures (Fung et al., 2008). Our results show that differential taxa between varieties are ubiquitous fungi in agricultural ecosystems, and some of these taxa are considered to be grapevine pathogens, for example *C. ramotenellum* and *Mycosphaerella tassiana* (Bensch et al., 2012), were more abundant in Pinot Noir musts (Fig. 1D).

Geographical signatures diminished during spontaneous wine fermentation as growth of fermentative yeasts reshaped the community diversity and composition, in particular *S*.*cerevisiae* (Fig. 2, S2; Supplementary Table S2, S3). Regardless of grape variety, fungal and yeast species diversity collapsed as alcoholic fermentation progressed, with a loss of the environmental fungi (Fig. 2A, S2A; Supplementary Table S3), indicating evolution through selection associated with wine fermentation. Non-*Saccharomyces* yeasts start the fermentation process (especially genera *Hanseniaspora, Candida, Pichia* and *Metschnikowia*), but they are quickly replaced by *S*.*cerevisiae* that lead fermentation until the end (Capozzi et al., 2015; Fleet, 2003). The inability of non-*Saccharomyces* yeasts to sustain their presence in ferments has been attributed to their oxidative and weakly fermentative metabolism, their sensitivity to increasing fermentation rate, ethanol and heat production induced by *S*.*cerevisiae* growth, and being less tolerant to low oxygen availability (Bokulich et al., 2016; Combina et al., 2005; Goddard, 2008). Thus, fermentation conditions, such as the chemical environment and interactions within the community drive the microbial pattern into a population dominated by *S. cerevisiae* (Fig. 2C, S2C; Supplementary Table S3) (Liu et al., 2017).

Spontaneous wine fermentation is characterised not only by significant intraspecific biodiversity (Cocolin et al., 2000), but also by high genetic polymorphism in the *S*.*cerevisiae* population present during the fermentation. Biogeography of *S*.*cerevisiae* has been previously reported from regional (Gayevskiy and Goddard, 2012; Knight and Goddard, 2015) and global (Legras et al., 2007; Liti et al., 2009) scales to small scales between vineyards within a single region (Börlin et al., 2016 ; Schuller and Casal, 2007), while Knight et al. (2019) demonstrated no vineyard demarcation between *S. cerevisiae* that could be ascribed to gene flow. Here, we show that distinct geographical differentiation between wine estates, with increased genetic divergence with distance (Schuller and Casal, 2007). A certain degree of mixed strains from various vineyard sites indicates gene flow among populations at small scales. Within the wine estates, the vineyards are within a 5 km radius from one another, and thus insect vectors like honeybees, wasps, and fruit flies, as well as birds, may homogenise the yeast populations (Francesca et al., 2012; Goddard et al., 2010; Lam and Howell, 2015; Stefanini et al., 2012). Shared staff and agricultural implements may also facilitate the movement and exchanges of *S*.*cerevisiae* populations between vineyards managed by the same estate (Goddard et al., 2010). Harvested grapes from various vineyards within the estate were processed at the same winery, where equipment surfaces may harbour large populations of *S*.*cerevisiae* and other yeasts under normal cleaning conditions, acting as potential reservoir for wine microbiota during spontaneous fermentation (Bokulich et al., 2013; Ciani et al., 2004; Sabate et al., 2002). All these factors contribute to the estate-specific pattern of *S*.*cerevisiae* distribution.

### 4.2 Association between fungal microbiota and composition of the wine metabolome

Previous research suggested that fungal microbiota structured and distinguished vineyard ecosystems impacting the aroma and quality of wine at the regional scale (Liu et al., 2020). Here we show that the geographical diversification observed in fungal communities, yeasts, and *S*.*cerevisiae* could translate to aromatic differences in wines within a single region. Wine aroma profiles highly associated with fermentative yeasts, in particular, *S*.*cerevisiae* displayed the most important effects on wine characteristics at this scale (Fig. 5, 6). Other fungal species, although non-fermentative (or not known to be associated) with wine fermentation, could produce some sensory-active compounds associated with wine aroma formation (Verginer et al., 2010), or substantially modulate vine health, growth, and fruit quality in the vineyard (Berg et al., 2014; Gilbert et al., 2014). The most widespread fungi *A. pullulans* are known to have an antagonistic effect on mould development, like *Botrytis cinerea*, causing grey rot, and *Aspergillus* spp., producing ochratoxin (Barata et al., 2012). Further, host-microbe interactions could promote plant resistance to environmental/abiotic stress thus benefiting crop production (Berg, 2009; Lugtenberg and Kamilova, 2009).

In the resultant wines, most volatile compounds were esters, alcohols, acids and aldehydes, many of which are fermentation-derived products. Some compounds were grape-derived, for example monoterpenes, and potentially modified by microbial metabolism during fermentation (Swiegers et al., 2005). Non-*Saccharomyces* yeasts dominating the initial spontaneous fermentation can contribute to the overall wine aroma profiles by producing flavour-active compounds, which depends on yeast species and strains (Capozzi et al., 2015; Jolly et al., 2014). Here, our work shows the presence of *T. delbrueckii, M. guilliermondii, and Hanseniaspora spp*. correlated with some volatile compounds, such as higher alcohols, ethyl esters, and acids (Fig. 5), thus potentially affecting wine characteristics. *M. guilliermondii*, which is known to produce *β*-glucosidases to release bound terpenoids (Silva et al., 2005), associated with monoterpenes including linalool, terpinen-4-ol, nerol, and *α*-terpineol, thus enhancing varietal aroma in wine and local *terroir* expression. Beyond fermentative contributions, the presence of non-*Saccharomyces* yeasts and the species diversity indirectly affect wine ecosystem function by altering the ecological dominance of *S. cerevisiae* through antagonistic interactions in early succession (Bagheri et al., 2017; Boynton and Greig, 2016). *S. cerevisiae* eventually dominate the fermentation, where they are naturally initially rare, by modifying the environment through fermentation, and drive wine ecosystem function (Boynton and Greig, 2016; Goddard, 2008). *S. cerevisiae* show genetic diversity at strain level and it is well documented that different genotypes produce variable amounts of volatile compounds as fermentation by-products, with desirable or undesirable impacts on wine aroma and flavour (Capece et al., 2012; Howell et al., 2004; Pretorius, 2000; Romano et al., 2003). Knight et al. (2015)found that *S. cerevisiae* genotypes and wine phenotypes were correlated with geographic dispersion, and regional populations produced distinct aroma profiles in Sauvignon Blanc wine fermentation. Our results further suggest that within a single region, wine aroma profiles were affected by the geographical origin and genomics of *S. cerevisiae* natural strains. As the keystone driver of the wine ecosystem, *S. cerevisiae* was associated with many ethyl esters and acetate esters, exerting the most powerful influences of multiple layers of fungal microbiota on wine quality and style (Fig. 5, 6). A causative relationship between the geographically differentiated fungi, yeasts, and *S. cerevisiae* should be established to show the impact on wine compounds for regional differentiation based on sensory characteristics.

## 5. Conclusions

Our study describes the diversity of fungal communities during spontaneous wine fermentation in carefully selected vineyards comprising two cultivars. As predicted, we observed ecological dominance *Saccharomyces* spp., but showed that geographical diversification is evident in the initial fungal community composition and the strain level diversity of *S. cerevisiae*. Fungal species correlated with wine volatile compounds, of which *S. cerevisiae* is likely the primary driver of wine aroma and characteristics within the sub-region (less than 12 km). A better understanding of how multiple layers of fungal microbiota vary at different scales, and the effects of these communities on agricultural ecosystems, provides perspectives for sustainable management practices maintaining the biodiversity and functioning thus optimising plant food and beverage production.

## Acknowledgements

We give our sincere thanks to the vignerons who kindly enabled sampling and provided wine samples. DL acknowledges support from a Ph.D. scholarship and funding from Wine Australia (AGW Ph1602) and a Melbourne Research Scholarship from the University of Melbourne. We would also like to acknowledge Master student Haoran Liu at the University of Melbourne for assistance in sample collection and yeast culture experiments.

## Figures and figure legends

**Fig. S1.**
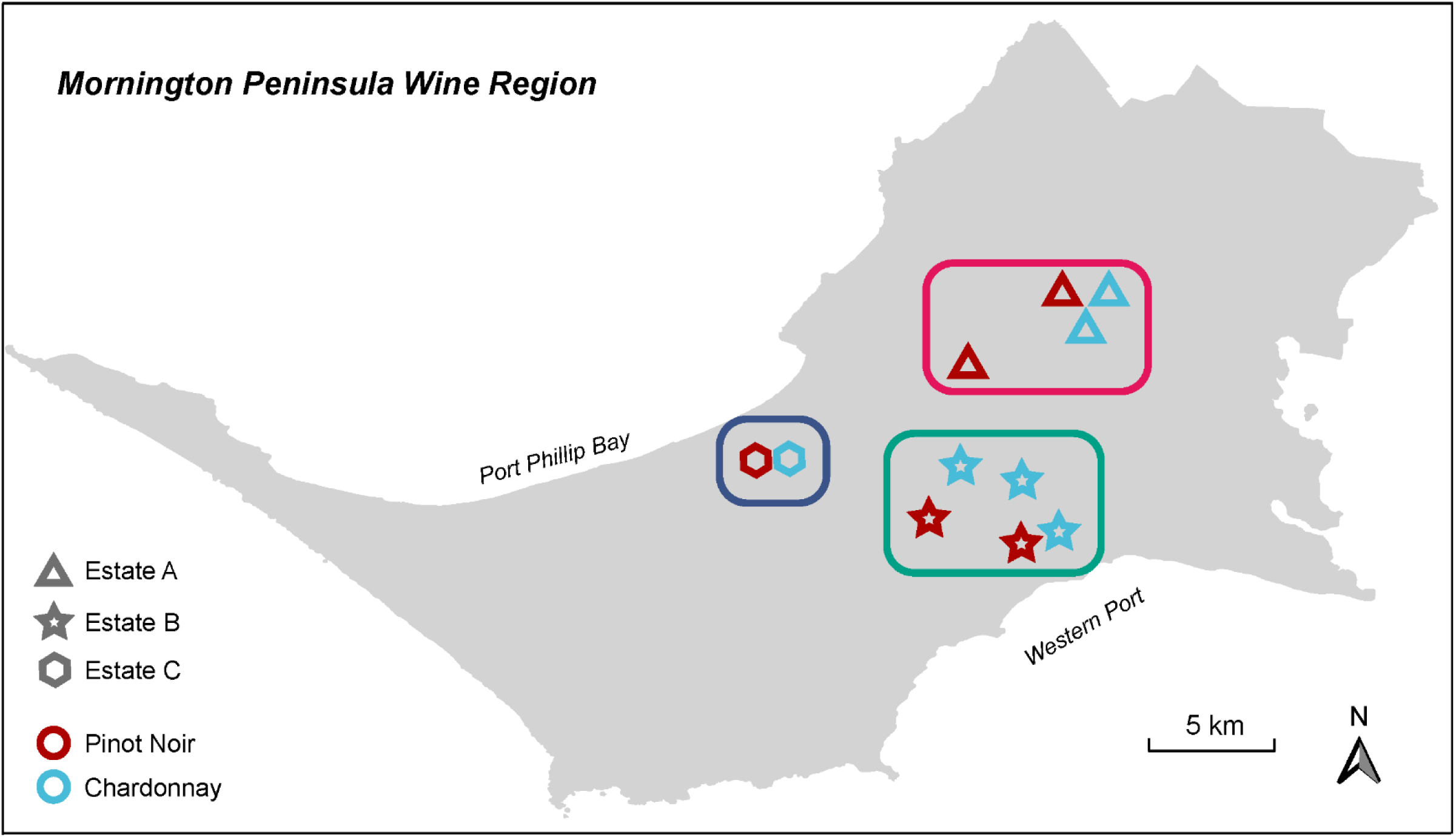
Sampling map of 11 vineyards from three wine estates in Mornington Peninsula wine region, Australia.

**Fig. S2.**
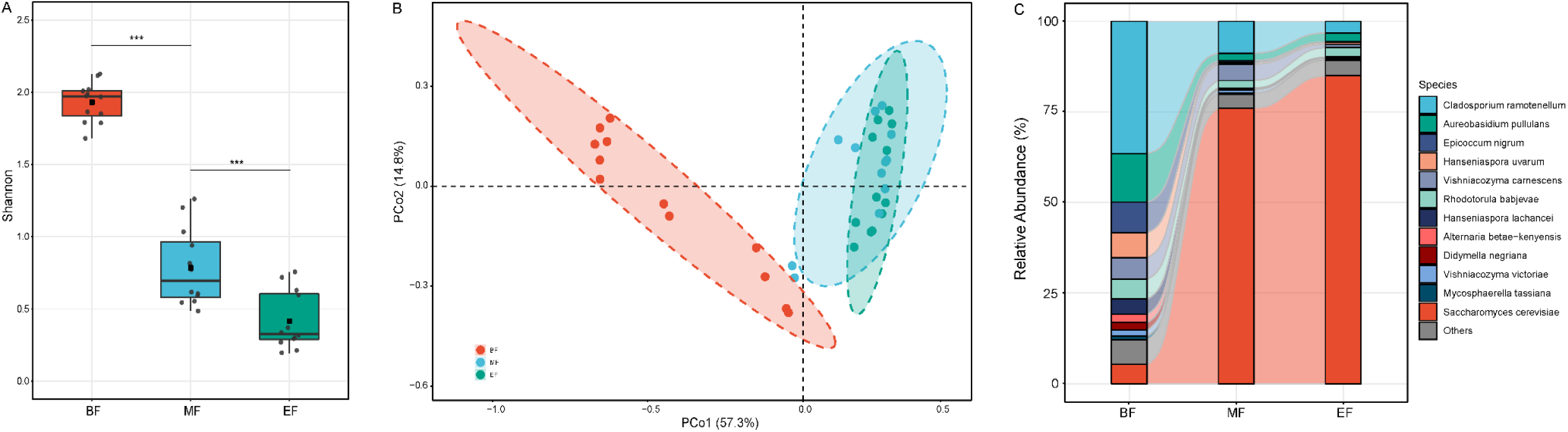
Stage of fermentation influences microbial diversity and composition of Chardonnay. (A) *α*-diversity (Shannon index) significantly decreases (*p* < 0.001***) during the fermentation; (B) Bray-Curtis distance PCoA of fungal communities according to the fermentation stage (BF, before fermentation; MF, at the middle of fermentation; EF, at the end of fermentation); (C) Relative abundance changes of major fungal species (12 species; relative abundance > 1.00%) that significantly differed by fermentation stage (ANOVA; FDR-corrected *p* value < 0.05).

